# Diversity and adaptability of RNA viruses in rice planthopper, *Laodelphax striatellus*

**DOI:** 10.1101/2023.10.08.561400

**Authors:** Qian-Zhuo Mao, Zhuang-Xin Ye, Jing-Na Yuan, Chao Ning, Meng-Nan Chen, Yu-Hua Qi, Yan Zhang, Ting Li, Yu-Juan He, Gang Lu, Hai-Jian Huang, Chuan-Xi Zhang, Jian-Ping Chen, Jun-Min Li

## Abstract

Although a large number of insect-specific viruses (ISVs) have recently been discovered from hematophagous insect, studies on ISV diversity and their association with phytophagous insect hosts were still insufficient. A systematic RNA virome investigation was performed for an important plant virus vector, small brown planthopper (SBPH), *Laodelphax striatellus*. A total number of 22 RNA viruses (including 17 novel viruses) belonging to various families were successfully identified and characterized. Subsequent analysis indicated that the overall RNA virus transcripts per million (TPM) in SBPH was relatively consistent throughout various different developmental stages of the insects, although the titers of individual viruses differ among different insect tissues, suggesting a delicate balance between ISVs and insect hosts. Moreover, analysis of virus-derived small interfering RNA (siRNA) demonstrated that siRNA mediated antiviral immune response of SBPH was activated in response to the replication of the discovered RNA viruses. Additionally, evaluation for potential cross-species ability of SBPH-ISVs showed that certain SBPH ISVs could successfully infect and replicate in another two rice planthoppers, the brown planthopper and the white-backed planthopper, through microinjection. In conclusion, the RNA virome and their adaptability in SBPH revealed in this study will contribute to a better understanding of the intimate relationship between ISVs and the host insects.

## Introduction

The next-generation sequencing technologies have revolutionized the identification of viral sequences in the past decade, redefining our understanding of viruses, especially for RNA viruses. The discovery of a large number of viral sequences has led us to realize that the presence of viruses has great ecological and evolutionary significance in the ecosystem (Dance, 2021; Koonin et al., 2022; Shi et al., 2016). It has been demonstrated that a number of insects can act as virus vectors for animals or plants, and the viruses within the insect vectors primarily consist of insect specific viruses (ISVs), vector-borne pathogenic viruses, as well as viruses from insect symbiotic microbes or digest materials (Bonning, 2019; Nouri et al., 2018). Complex interactions exist between multiple viruses, viruses-host antiviral pathways, and viruses with other microorganisms in insect vectors, which may affect the physiology and vector competence of insects (Nouri et al., 2018; Olmo et al., 2023; Patterson et al., 2020). Therefore, virome characterization of insect vectors is crucial to gain a better understanding of the co-evolution and interaction between viruses and insects.

The virome composition of insects is related to host species, geographic locations, seasons and environmental factors (de Almeida et al., 2021; French et al., 2023; Huang et al., 2021; N. Li et al., 2023; Y. Xu et al., 2022). Research on the mosquito virome, an important vector of human and animal infections, revealed that viral abundance and composition varied seasonally and with mosquito species (de Almeida et al., 2021; Feng et al., 2022). The diversity of RNA virome in the agricultural important plant virus vector whitefly was associated with host cryptic species, and some viruses were capable of breaking the cryptic species barrier through micro-injection (Huang et al., 2021). Furthermore, at the level of individual viruses, many ISVs have been identified and demonstrated to have an effect on insect physiology or pathogen transmission (Jia et al., 2021; Patterson et al., 2020; Wang et al., 2017;). Previous study showed that pathogenic ISVs, such as baculoviruses, can kill hosts and be used as bio-pesticides (Lacey et al., 2015). Recent years, more and more ISVs are discovered and exhibit various adaptive interactions with insect host. Some partiti-like viruses in armyworms can increase the host’s resistance to pathogenic virus despite harming the host’s growth and reproduction, exhibiting conditional mutualistic symbiotic relationships with hosts (P. Xu et al., 2020). In mosquitoes, several ISVs influence the replication or spread of arboviruses such as DENV and ZIKV in the insect body, and modulate the vector competence for dengue (Baidaliuk et al., 2019; Goenaga et al., 2015; Öhlund et al., 2019; Olmo et al., 2023). Insects are generally co-infested by multiple virus species, thus competition for survival resources and synergistic interactions were commonly observed among various viruses and between viruses and its hosts. Therefore, the virome diversity may reflect the adaptability of insects to various survival factors, and contribute to the research on the dynamic nature of virus incidence within its hosts.

The small brown planthopper (SBPH), *Laodelphax striatellus* (Delphacidae; Hemiptera), is a sap-sucking pest that feeds primarily on rice, and transmits a variety of important crop viruses (Cao et al., 2018; J. Li et al., 2013). In addition to SBPH-borne crop viruses, several ISVs in the taxon of *Dicistroviridae*, *Iflaviridae* or *Fijivirus* were recently been reported or submitted to GenBank (GuY et al., 1988; Lu et al., 2022a; Wu et al., 2019). Nevertheless, the overall virome of SBPH is still not available and has not been characterized. In this study, we conducted a systematic identification and analysis of the entire RNA virome in SBPH, identified 22 viruses in the SBPH, 17 of which were novel viruses. Moreover, we examined the infestation characteristic of RNA viruses in SBPH tissues and their adaptation to SBPH development and the small RNA antiviral pathway. Considering the close relationship of SBPH with other two planthopper species, the brown planthopper (BPH) *Nilaparvata lugens* and the white-backed planthopper (WBPH) *Sogatella furcifera*, the ability of ISVs from SBPH to cross-infect other two planthopper species were also investigated. This study will contribute to gain a better understanding of adaptation and coevolution between viruses and hosts.

## RESULTS

### Datasets used for SBPH virome analysis

A total of 39 transcriptome datasets of SBPH were assembled/reassembled and the N50 of each library (assembles with Trinity) was calculated. Information related to these datasets was provided in Supplementary Table 1. Among them, 28 datasets were obtained from the NCBI SRA repository that were submitted by seven different universities or institutions in China (Supplementary Figure S1 and Supplementary Table S1). Additionally, we sequenced two lab-reared samples that were maintained in NBU. Moreover, we obtained six field samples of SBPH from various locations in China (Hangzhou, Jurong, Xinxiang, Xinzhou, Dalian, and Shenyang) and RNA-seq libraries were subsequently constructed (Supplementary Figure S1 and Supplementary Table S1). All of the SBPH datasets were subsequently used for the RNA virome analysis.

### Diversity of RNA viruses identified in SBPH

In all of the assembled libraries, a total number of 22 RNA viruses were identified, of which 17 are novel ISVs (Supplementary Table 2). The five known ISVs included three iflaviruses (Laodelphax striatellus iflavirus 1 (LSIV1), Laodelphax striatellus picorna-like virus 2 (LSPV2), Laodelphax striatella honeydew virus 1 (LSHV1)), one triatovirus (Himetobi P virus (HiPV)), and one fijivirus (Laodelphax striatellus reovirus (LSRV)) (GuY et al., 1988; Lu et al., 2022). Based on the viral genome organization and phylogenetical analysis, the 17 newly discovered viruses were classified to 13 different viral families or orders. These include two dsRNA viruses (*Fusariviridae* and *Partitiviridae*), 11 +ssRNA viruses (*Permutotetraviridae*, *Flaviviridae*, *Botourmiaviridae*, *Iflaviridae*, *Solemoviridae* and *Narnaviridae*) and four -ssRNA viruses (*Bunyavirales*, *Phenuiviridae* and *Aliusviridae*). The conserved domains of the viral genome of these 17 newly discovered viruses, along with the corresponding closely related homologous viruses, are presented in Figure 1. The taxonomical status of these viruses were inferred with phylogenetic trees based on the predicted viral RdRP protein sequences (Figure 2).

**Figure 1.**
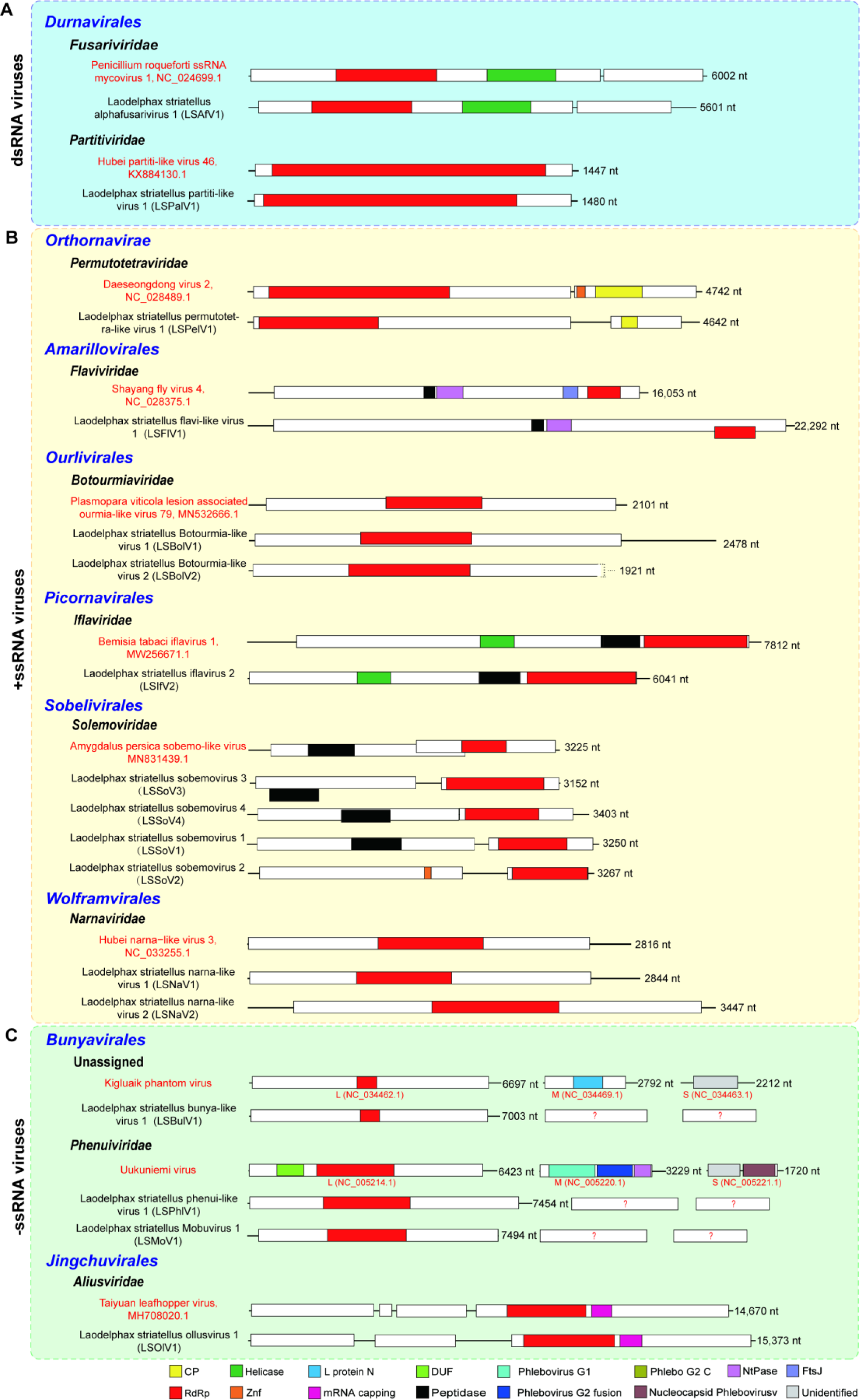
Genomic structures of novel viruses identified in SBPH. The viruses were taxonomically classified into 3 groups, including dsRNA viruses (A), +ssRNA viruses (B) and –ssRNA viruses (C). Viruses are arranged by order and family, and in the group of viruses belonging to the same family, a phylogenetically close reference virus is listed on the top with red font, followed by novel viruses. The Viral proteins are colored according to their putative functions. GenBank accession number are provided following the name of the reference viruses.

**Figure 2.**
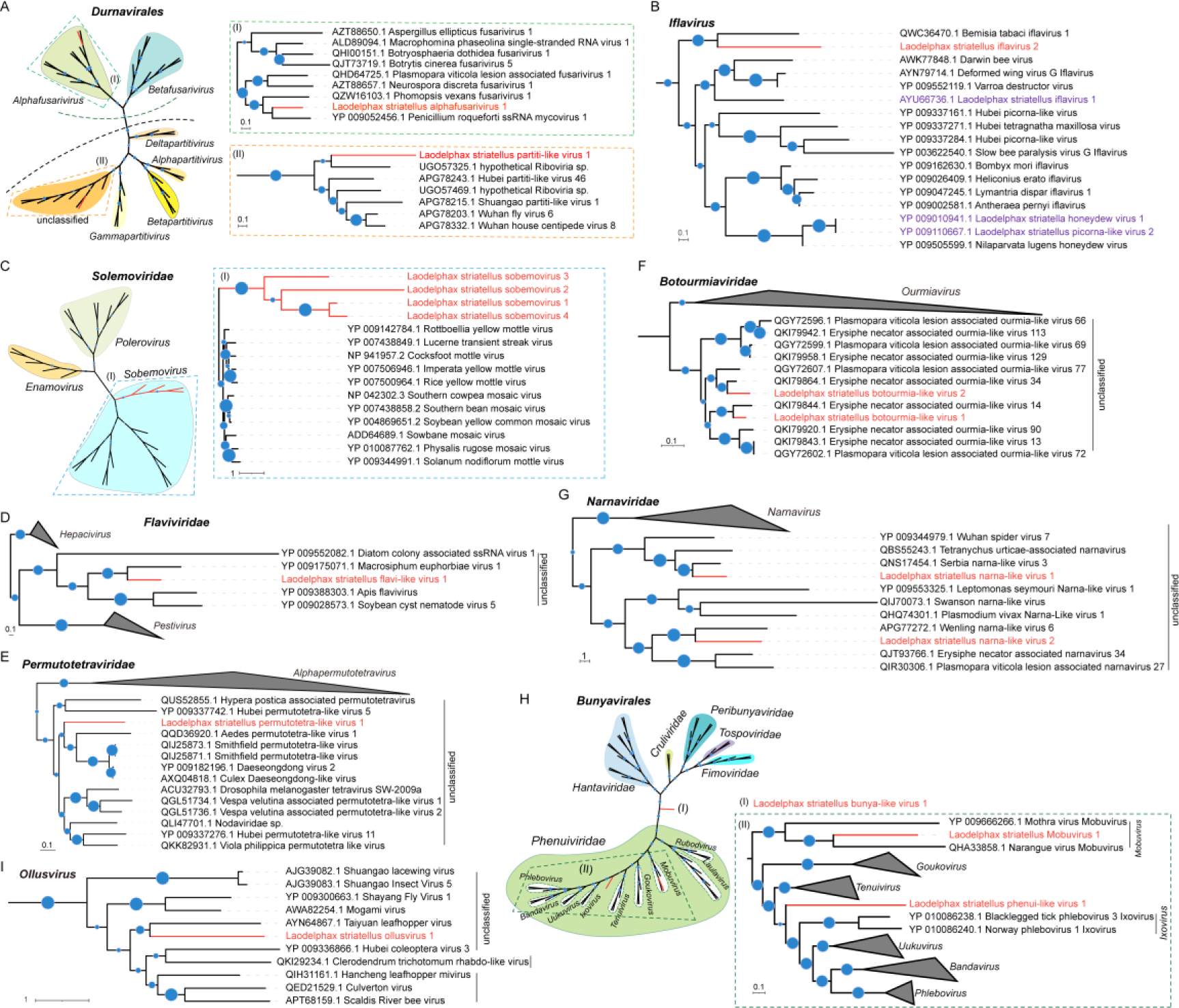
Phylogenetic trees of the novel RNA viruses identified in SBPH. Trees for *Durnavirales* (A), *Iflavirus* (B), *Solemoviridae* (C), *Flaviviridae* (D), *Permutotetraviridae* (E), *Botourmiaviridae* (F), *Narnaviridae* (G), *Bunyavirales* (H), and *Aliusviridae* (I) are based on the maximum likelihood method and inferred from conserved viral RdRP domains. Novel viruses are shown in red font. Nodes with bootstrap values >50% are marked with blue circles, and the larger circles indicated higher bootstrap values. In panels A, C and H, taxonomic overview of viruses at order or family level are shown on the left, and a close-up view of the viruses are shown in the boxes with the color of dotted frames corresponding to the interested clusters. The viral sequences used in this study were extracted from GenBank, the accession numbers and other related details are listed in Supplementary Table 3.

#### dsRNA viruses

Three novel dsRNA viruses were identified from the transcriptomes of SBPH. Each virus belongs to a different family, including *Fusariviridae*, *Partitiviridae*, and *Spinareoviridae* (Figure 1A, Supplementary Table 2). Phylogenetically, the novel virus Laodelphax striatellus alphafusarivirus 1 (LSAfV1) was closely related to fungal viruses (Figure 2A-I) (Gong et al., 2021). On the other hand, Laodelphax striatellus partiti-like virus 1 (LSPalV1) clustered together with viruses reported or found in invertebrate hosts (Figure 2A-II) (Shi et al., 2016). Notably, the dsRNA viruses were predominantly found in laboratory samples, while only the virus from the *Partitiviridae* was detected in field samples from Shenyang and Dalian (Figure 3).

**Figure 3.**
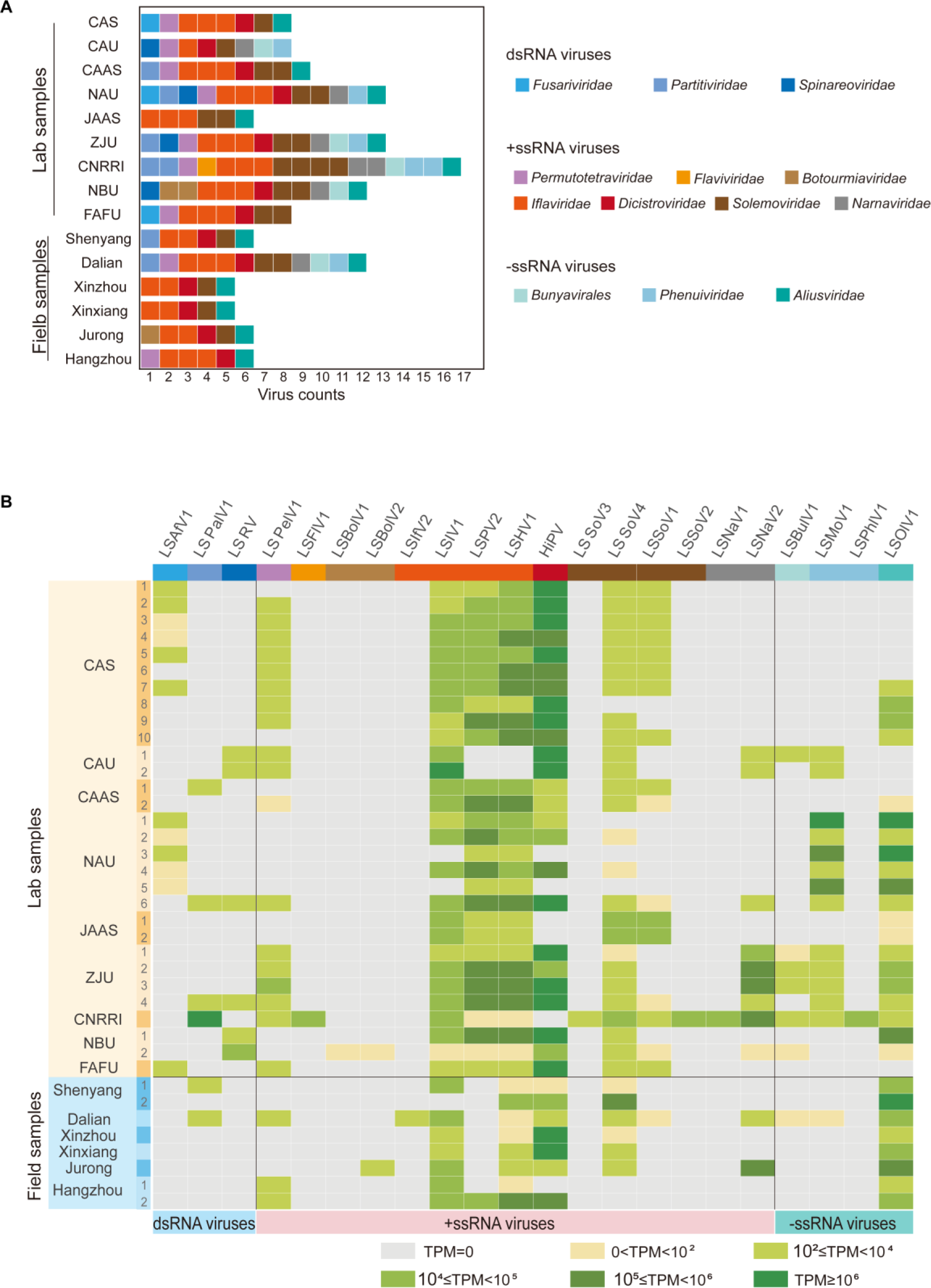
RNA virome composition and abundance across different datasets. (A) RNA virome composition of different lab and field samples; (B) Virus distribution across different databases. Different colors were used to indicate the abundance ranges determined by transcripts per million (TPM).

#### +ssRNA viruses

The +ssRNA viruses are the dominant type of SBPH RNA viruses in terms of both viral diversity and abundance. A total of 15 +ssRNA viruses were identified in SBPH, including 11 novel viruses and 4 reported viruses, which could be assigned to 7 families (Supplementary Table 2, Figure 1B).

A novel iflavirus, Laodelphax striatellus iflavirus 2 (LSIfV2), belonging to the family *Iflaviridae*, was discovered in SBPH based on its genome structure and phylogenetic analysis (Figure 1B and Figure 2B). Previous studies showed that a number of picorna-like viruses (mainly belonging to the families of *Iflaviridae* and *Dicistroviridae*), such as HiPV, LSIV1, LSHV, and LSPV, commonly infect SBPH and comprised the major part of SBPH RNA virome (Supplementary Table S2) (Gu et al., 1988; Wu et al., 2019). Our work also indicated that these viruses are the important core viruses prevalent in various laboratory and field samples with high abundance (Figure 3).

Interestingly, four novel viruses were identified and assigned to the family of *Solemoviridae* (Figure 1B and Figure 2C), a plant/invertebrate-associated viral family (Shi et al., 2016; Sõmera et al., 2021). These solemoviruses distributed in all laboratories SBPH cultures and the majority of field samples (Figure 3).

A novel permutotetra-like virus (LSPelV1) was found in samples from seven laboratories and two field-sampling locations, while another novel flavi-like virus (LSFlV1) was discovered only in lab samples submitted by CNRRI (Figure 3). Both of these viruses are phylogenetically related to insect-specific viruses (Figure 2D and E).

Four novel viruses belonging to the family of *Botourmiaviridae* and *Narnaviridae* were identified and phylogenetically related to fungal viruses (Figure 1B, Figure 2F and G). The two botourmia-like viruses were found only in the NBU lab sample with low abundance, whereas the two narna-like viruses were presented in SBPH samples derived from 5 laboratories and 2 field sampling locations (Figure 3). Considering that a number of narnaviruses were previously reported in arthropods along with the detection of fungal sequences (Chandler et al., 2015), this viral clade may be originated from fungi that colonized insects.

#### -ssRNA viruses

Four novel -ssRNA viruses were successfully identified from various SBPH datasets. These viruses included three viruses of the order *Bunyavirales* and one virus in the order *Jingchuvirales* (Figure 1C, Figure 2H, I and Supplementary Table 2). The bunya-like viruses were detected in samples from five labs and one field sample (Figure 3). Among them, the Laodelphax striatellus bunya-like virus 1 (LSBulV1) was phylogenetically separated from the known family while the two viruses were clustered with ISVs in the family of *Phenuiviridae* (Figure 2H). The *Jingchuvirales*, a new viral order established in 2022, comprised of viruses infecting arthropods that were globally distributed (Di Paola et al., 2022). A novel ollusvirus, Laodelphax striatellus ollusvirus 1 (LSOlV1), discovered in this study exhibited as the most prevalent -ssRNA virus across the samples derived from six labs and all six field locations (Figure 2I and Figure 3).

### Distribution and abundance of SBPH viruses in different tissues and genders

The relative abundance of RNA viruses in the guts (Gut), salivary glands (SG), fat bodies (FB), female reproductive systems (FR), male reproductive systems (MR), and residues (Re) of SBPH was determined through comparative transcriptome analysis. Our results demonstrated that 11 viruses were presented in our lab populations, with varying abundance across tissues. SG and Re exhibited the highest virus abundance, while the FR showed the lowest levels (Figure 4A). Among these viruses, three iflaviruses (LSIV1, LSPV2, and LSHV1) accounted for over 65% of the total viral abundance in each tissue. The narna-like virus (LSNaV2) accounted for 11% and 30% of the total viral abundance in SG and MR, respectively, while the jingchu-like virus (LSOlV1) accounted for 11% in FR (Figure 4A). Additionally, LSNaV2 was much more abundant in the Gut and MR compared to other tissues. LSSoV1 was not detected in the guts or reproductive systems (FR and MR). LSBulV1 was nearly absent in the Gut and FB, whereas LSBulV1, LSOlV1, and LSPhlV1 were much more abundant in FR compare to other tissues (Figure 4B). The abundance of other viruses, including LSPelV1, LSSoV4, LSIV1, LSPV2, and LSHV1, was comparable in different tissues (Figure 4B).Regarding the total viral abundance, no significant differences were observed between female and male adult SBPH (Figure 4C). However, for the individual viruses, LSPelV1 showed higher abundance in males than that of the females (Figure 4D). Nonetheless, there were no significant differences in the distribution of this virus among female and male reproductive systems. Conversely, two bunya-like viruses, LSBulV and LSPhlV1, exhibited significantly higher abundance in females compared to males, consistent with their elevated expression in FR compared to MR (Figure 4B and D). Moreover, no significant differences were observed for the abundance of other viruses between males and females (Figure 4D).

**Figure 4.**
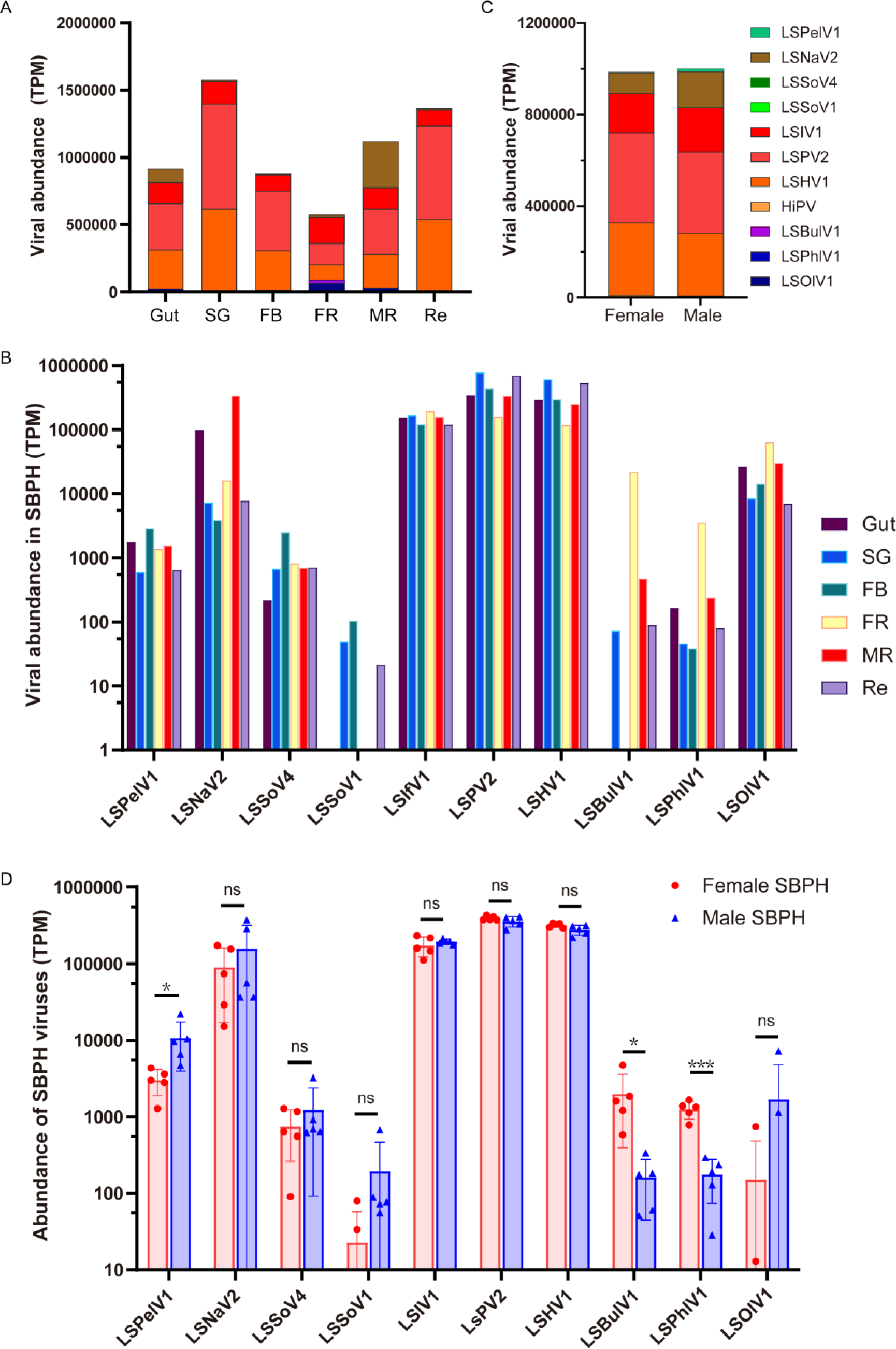
Distribution and abundance of RNA viruses in different tissues and genders of SBPH. (A) Total abundance of viruses in guts (Gut), salivary glands (SG), fat bodies (FB), female reproductive systems (FR), male reproductive systems (MR), and residues (Re) of SBPH; (**B**) abundance of individual viruses in different tissues of SBPH; (C, D) Comparison of virome abundance (C) and abundance of individual viruses (D) between male and female adults.

### Virome diversity and abundance in different developmental stages of SBPH

The viral abundance dynamics of individual viruses in different development stages (from egg to adult) were evaluated for the SBPH population in our laboratory. The presence of 11 viruses was revealed by transcript mapping analysis. The results suggested that the abundance of total RNA viruses in SBPH was relatively stable at a level of 1.0×10^6^ TPM throughout the different development stages (egg, nymph, and adult stages) (Figure 5A).

**Figure 5.**
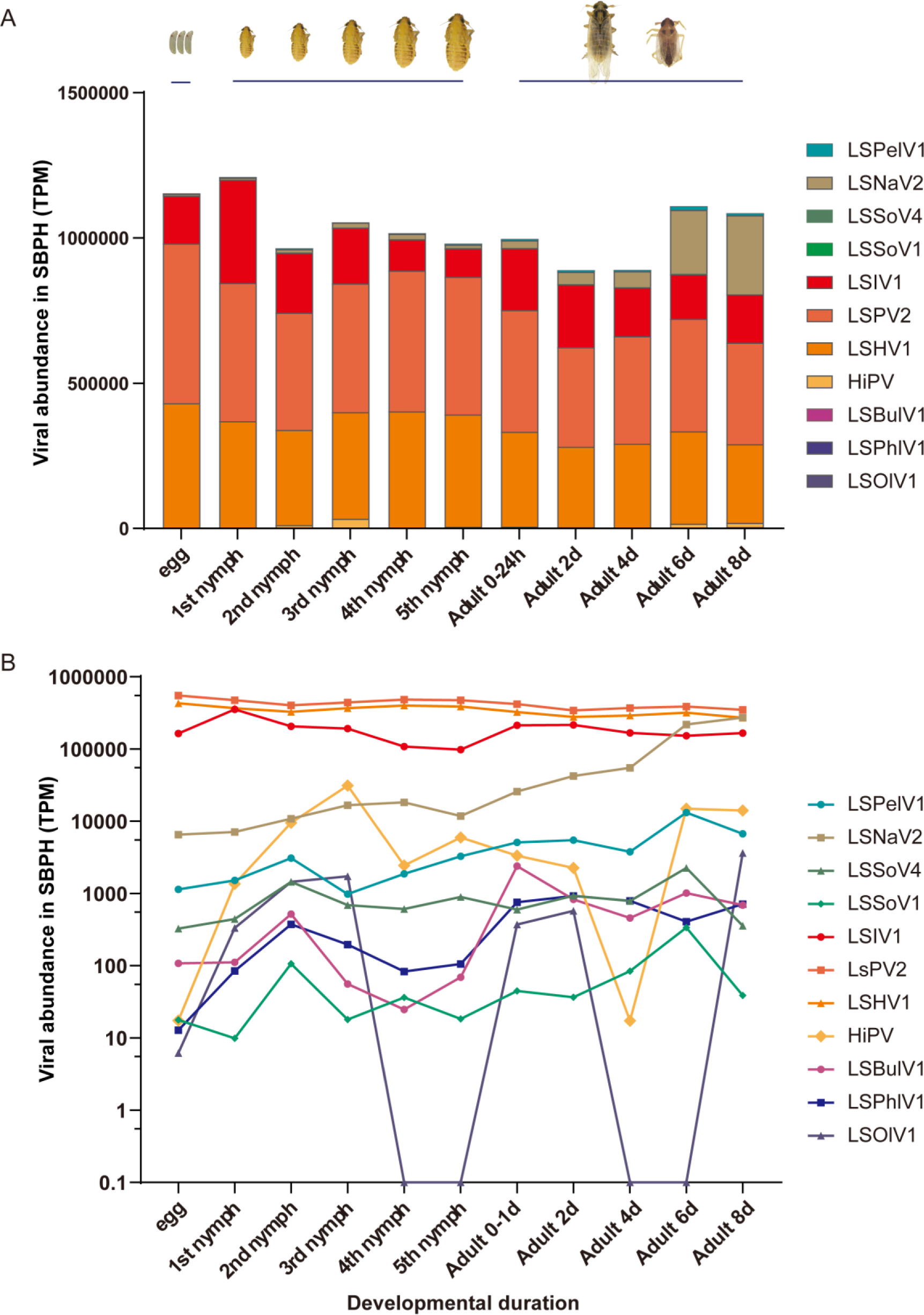
Abundance of RNA viruses in different development stages of SBPH lab populations. (A) The abundance of total RNA viruses throughout different development stages of SBPH, including eggs, 1st-5th nymphs, and adults 0-1 d, 2 d, 4 d, 6 d, and 8 d post eclosion; (B) abundance of individual viruses throughout different development stages of SBPH.

For individual viruses, most of them exhibited consistent infestation levels throughout the various developmental stages except for HiPV and LSOlV1 (Figure 5B). The abundance of each virus species maintained respective levels of viral abundance throughout the life cycle of SBPH (Figure 5B). These findings imply the existence of viral load thresholds for the proliferation of different viruses in SBPH, enabling persistent virus infestation in the insects. Notably, two closely related iflaviruses, LSPV2 and LSHV1, exhibited almost identical infestation levels and proliferation patterns (Figure 5B). In contrast, HiPV, a virus frequently observed in various SBPH databases and characterized by consistently high levels of abundance (Chen et al., 2015; J. Li et al., 2014), displayed an unstable viral expression pattern in our laboratory population (Figure 5B). The reasons behind this pattern could be either a low viral infection rate in this population or the regulation of persistent SBPH infestation by this virus through several unknown factors. As for the narna-like virus LSNalV2, its viral load increased significantly with the development of SBPH, particularly after eclosion of the adults (Figure 5B). Considering its taxonomic relation to fungal viruses, the infection pattern of this virus may be associated with the function of its fungal host (Hillman & Cai, 2013).

During the egg stage of the insects, each virus was detected at different abundant levels (Figure 5B). Of these, HiPV has been consistently reported to be incapable of vertical transmission in host insects (Toriyama et al., 1992). However, the transcriptome analysis demonstrated its presence at low abundance, possibly due to the lack of surface washing of the tested egg samples, thus allowing the detection of the virus on the egg surface. Consequently, it remains unclear whether viruses with lower viral loads in eggs (such as LSSoV1, LSPhlV1, and LSOlV1) are vertically transmitted in the SBPH population. Contrarily, several SBPH viruses (LSBulV1, LSSoV4, LSPelV1, LSNalV2, LSIV1, LsPV2, and LSHV1) exhibited higher viral loads in eggs, implying that they might be vertical transmitted (Figure 5B).

### Analysis of virus derived siRNAs in SBPH lab populations

Viral replication activates RNA interference (RNAi) pathways mediated by small interfering RNA (siRNA), so vsiRNA analysis is essential for detecting RNA viruses and obtaining putative viral sequences that lack detectable sequence similarity to known viruses (Gammon & Mello, 2015; Webster et al., 2015). To investigate the infestation of SBPH-specific viruses in insects and host siRNA-mediated immune regulation, we conducted both transcript and small RNA sequencing on our SBPH laboratory populations. Among 22 RNA viruses identified in SBPH, siRNAs of this SBPH lab sample were mapped to eight viruses after filtering out host genome sequences. These included six +ssRNA viruses (LSIV1, LSPV2, LSHV1, HiPV, LSSoV4, and LSSoV1), and two -ssRNA viruses (LSBulV1 and LSOlV1) (Figure 6A). With the exception of HiPV and LSOlV, the virus-derived siRNAs (vsiRNAs) exhibited the typical size distribution and polarity pattern: predominantly distributed at 19 to 23 nucleotides, with a peak at 22 nt (Figure 6A). Furthermore, they originated nearly equally from both sense and antisense strands of the viral genomic RNA (Figure 6A). Both HiPV-vsiRNA and LSOlV-vsiRNA displayed a significantly positive-strand bias and a broad range of sizes peaking at 22 nt (Figure 6A). These findings indicate that the virus can replicate within SBPH and trigger the antiviral RNAi pathway. Additionally, an analysis of the vsiRNA distribution in the corresponding genomes/segments revealed widespread distribution but with noticeable asymmetric hotspots on both strands, suggesting that these regions might be preferentially targeted for cleavage by the host immune system (Figure 6A). We then calculated and compared the number of vsiRNA reads per transcript (referred to as vsiRNA ratio) (Webster et al., 2015). The results unveiled variations of the vsiRNA ratios for the different viruses. Notably, the two solemo-like viruses (LSSoV4 and LSSoV1) displayed above ten times higher vsiRNA ratios in comparison to the iflavirus LSIV1 (Figure 6B).

**Figure 6.**
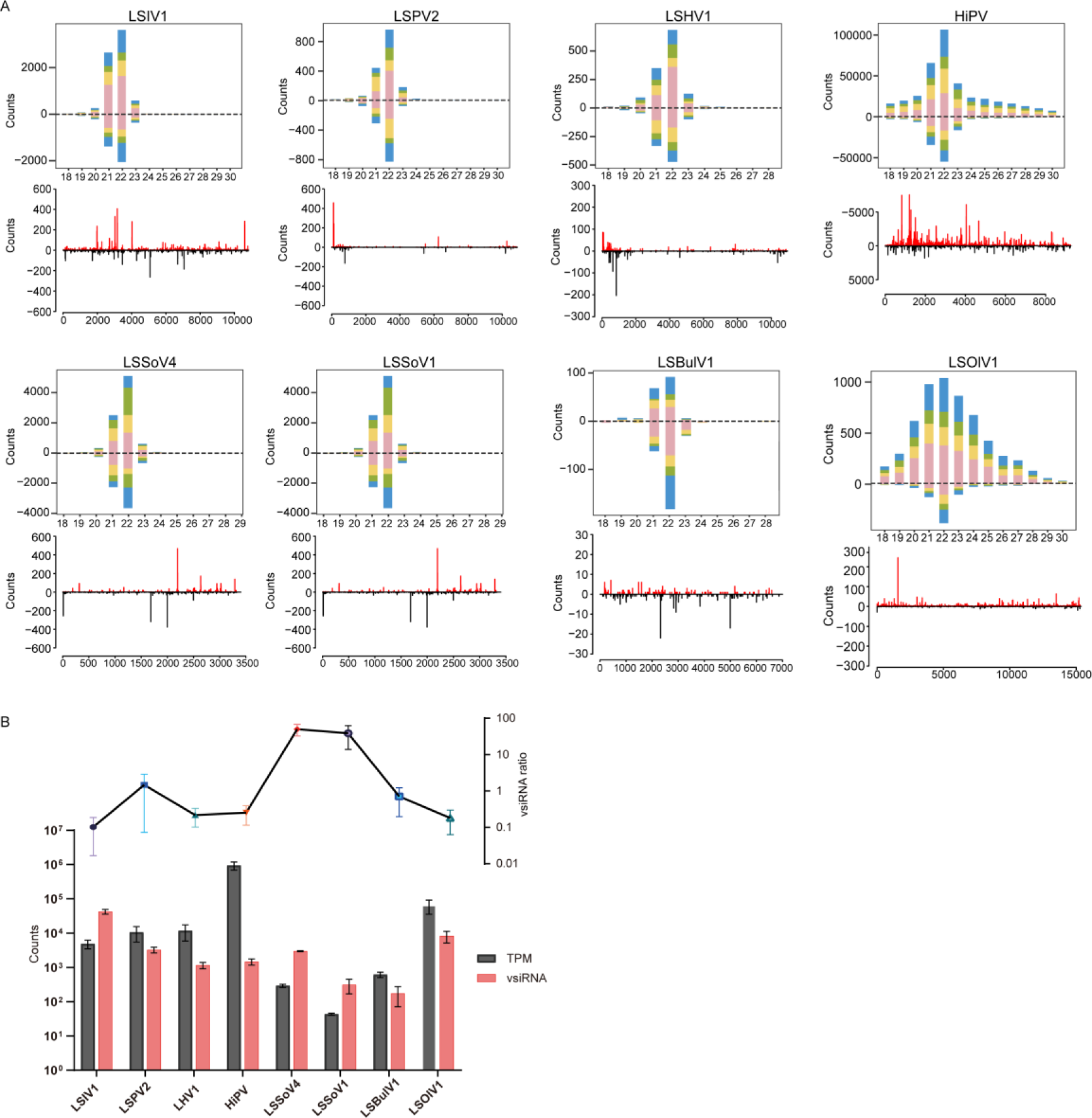
SBPH-ISV derived small interfering RNAs in SBPH lab-populations. (A) Profiles of vsiRNAs. The size distribution of vsiRNAs was shown in the upper panel, while the distribution of vsiRNAs in the corresponding strands was shown in the lower panel. (B) The vsiRNA ratio of different SBPH-ISVs. The total vsiRNA counts the TPM of each virus were showed in the column diagram, and the counts of vsiRNA reads produced per transcript of each ISV (vsiRNA ratio) were showed in the upper panel.

### Evaluation for cross-species ability of SBPH ISVs

Given the close phylogenetic relationship among three planthopper species (SBPH, BPH and WBPH) (Ammar & Nault, 2002), the potential infection and replication ability of seven SBPH-ISVs in WBPH and BPH was investigated. SBPH-ISV inoculum was injected separately into 3rd-instar individual nymphs of WBPH and BPH (Figure 7A) and the presence of the SBPH-ISVs in the inoculum was confirmed by RT-PCR (Figure 7B).

**Figure 7.**
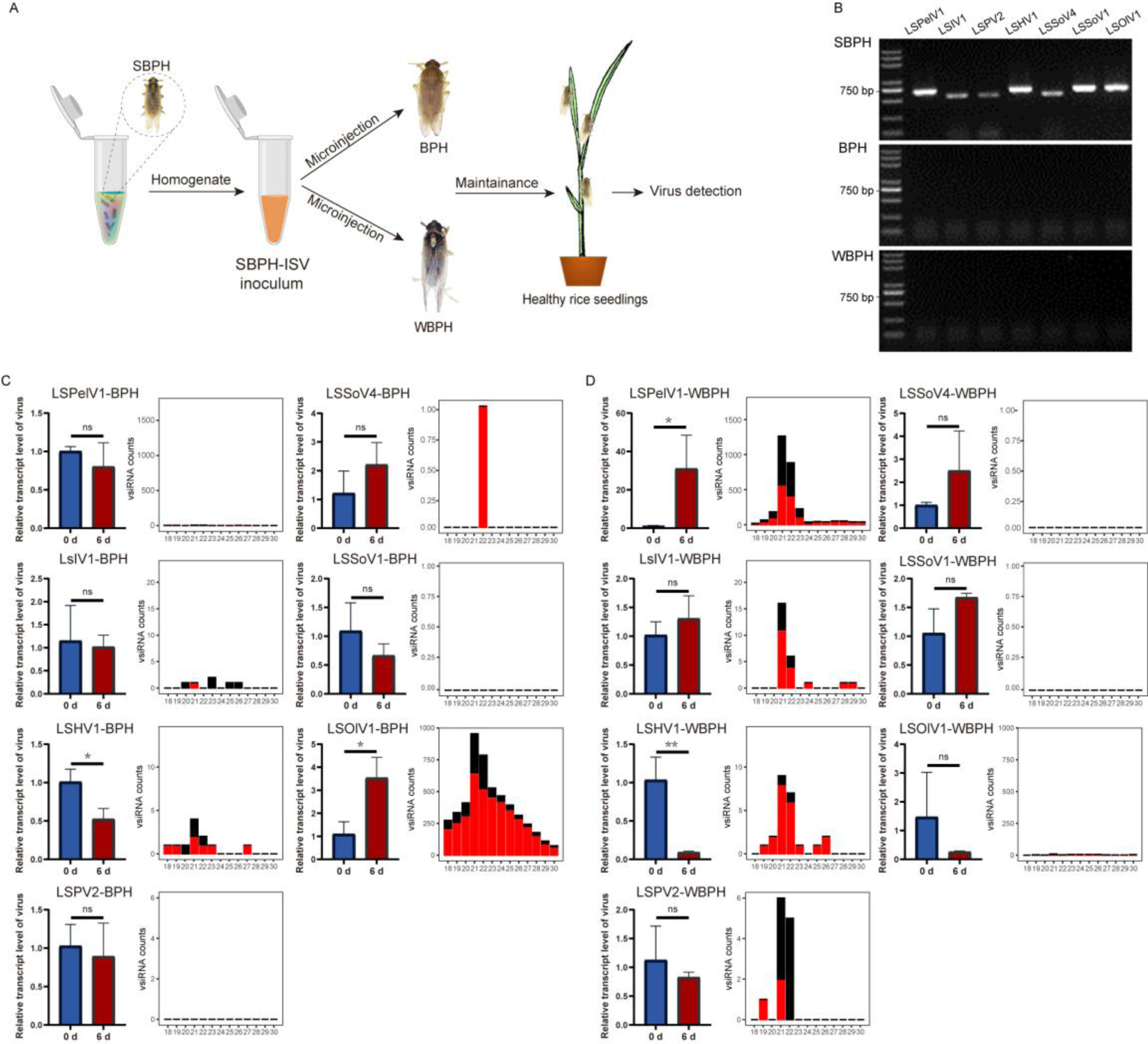
Evaluation of the ability of SBPH-ISVs to cross-infect other planthopper species WBPH and BPH. (A) Overall strategy to evaluate the cross-infection ability of SBPH-ISVs to other planthopper species WBPH and BPH. SBPH-ISV inoculum was prepared and injected separately into 3rd-instar nymphs of WBPH and BPH. (B) Detection of SBPH-ISVs in the SBPH-ISV inoculum and in the WBPH and BPH populations pre-injection. The presence of SBPH-ISVs in each sample was determined using RT-PCR. (C) Replication assessment of SBPH-ISVs in other planthopper species BPH and WBPH. Left column diagram showed the relative transcript level of SBPH-ISVs in BPH and WBPH at 0 and 6 DPI. A pool of 15 insects were collected for viral load detection using RT-qPCR, and 3 independent biological replicates were performed. Right diagram showed profiles of SBPH-ISV derived siRNAs in BPH and WBPH. The microinjected insects at 6 DPI were performed siRNA sequencing and analysis.

Among the seven SBPH-ISVs injected into BPH, LSOlV1 exhibited a significant increase in viral load at 6 days post-injection (DPI), accompanied by a high abundance of vsiRNA production (Figure 7C). In contrast, the viral load of LSHV1 decreased without vsiRNA induction (Figure 7C). The remaining viruses (LSPelV1, LSIV1, LSSoV4, LSSoV1, and LSPV2) did not display difference in viral loads or induce vsiRNA production (Figure 7C). For WBPH at 6 DPI post injection, LSPelV1 exhibited an increase in viral load along with a high abundance of vsiRNA production (Figure 7C). Similar to the results observed for BPH, the viral load of LSHV1 decreased without vsiRNA production. The other viruses also did not exhibit changes in viral loads or trigger vsiRNA production (Figure 7C). In conclusion, our findings suggest that certain SBPH-ISVs are capable of infecting and replicating in closely related planthopper species, such as LSOlV1 in BPH and LSPelV1 in WBPH.

## Discussion

In this study, we conducted a comprehensive and systematic analysis to identify and analyze RNA viruses in the planthopper SBPH. We successfully identified 22 RNA viruses, including 17 novel viruses from diverse families or orders (Supplementary Table 2). A number of identified RNA viruses were assigned in unclassified clades in the families of *Partitiviridae*, *Permutotetraviridae*, *Flaviviridae*, *Botourmiaviridae*, *Narnaviridae*, and *Phenuiviridae* (Figure 3), suggesting novel genera might be existed within these families. Additionally, we encountered one virus that could not be classified within any of the known viral families in the *Bunyavirales* order, implying that this virus might belong to a new family.

The SBPH viruses identified in our study included ISVs as well as viruses associated with plants or symbiont fungi. Most SBPH viruses clustered with invertebrate viruses from families of *Partitiviridae*, *Flaviviridae*, *Iflaviridae*, *Phenuiviridae*, *Dicistroviridae*, and *Aliusviridae* (Figure 3), indicating that they might specific to SBPH. Interestingly, considering that SBPH is an important plant virus vector, the four novel sobemoviruses discovered in our study (Figure 2C), as well as another fijivirus reported previously, exhibited close phylogenetic relationships with plant viruses (Lu et al., 2022a; Sõmera et al., 2021). This close relationship between insect vector and plant viruses strongly supports the hypothesis that plant viruses might be evolved from invertebrate viruses (Dolja et al., 2020). However, it remains unclear whether SBPH specific sobemoviruses pose a threat to crops or if SBPH can transmit viruses from the same taxon. Therefore, characterizing the SBPH virome is crucial for identifying potential plant viruses that can potentially endanger agricultural production. The second group of viruses identified in our study clustered with fungal viruses belonging to the families such as *Fusariviridae* and *Narnaviridae* (Gong et al., 2021; Hillman & Cai, 2013; Roossinck, 2019). These viruses were found abundantly in different populations (Figure 2C). The narna-like virus LSNaV2, in particular, exhibited high infestation rates in our laboratory populations and was distributed widely within various insect tissues (Figure 4 and 5). These findings suggest that the host fungi of LSNaV2 might be closely associated with SBPH. Thus, the virome, as a significant component of the insect microbiome, interacts with insects and symbiotic microbes in various ways.

Previous research has demonstrated that individual insects and insect groups can act as reservoirs for numerous viruses (Bonning, 2019; Öhlund et al., 2019; Olmo et al., 2019). Our experimental results revealed that while the titers of individual viruses differ among different insect tissues, the overall RNA virus transcripts per million (TPM) in whole insects remain relatively consistent throughout different developmental stages (Figure 4 and 5). The TPM value serves as an indicator of viral transcription and replication, and the consistent viral load of total RNA viruses suggests the presence of a delicate balance between viruses and insects (Aguiar et al., 2015, 2016; Gammon & Mello, 2015). This equilibrium implies that viruses can only utilize limited resources for persistent infection. To adapt to host selection pressures with limited resources, all viruses within insects establish synergistic or antagonistic relationships to ensure their survival and propagation among individuals and populations of insects (Bonning & Saleh, 2021).

In order to maintain the balance between persistent virus infection and insect survival, the innate antiviral immunity pathways of the host must play essential roles. In insects, RNA interference pathways detect virus infection and initiate an antiviral response to limit virus replication (Bonning & Saleh, 2021; Olmo et al., 2019). The production and abundance of vsiRNAs are involved in the processing of viral RNA products. The sRNA patterns from both SBPH-ISVs and SBPH-borne plant viruses display typical dicer-mediated degradation signatures, ranging from 18 to 25 nt in length and showing peak detection at 22 nt. This consistency in vsiRNA patterns between ISVs and vector-borne viruses implies that they were all regulated by dicer-mediated RNAi (Aguiar et al., 2016; Gammon & Mello, 2015; J. Li et al., 2013). Similar patterns of vsiRNAs have been observed in grasshoppers, thrips, and whiteflies (Chiapello et al., 2021; Huang et al., 2021; Y. Xu et al., 2022). Moreover, vsiRNA profiles for RNA viruses in other insects, such as mosquitoes, fruit flies, and leafhoppers, primarily peak at 21 nt (Aguiar et al., 2015, 2016; Lan et al., 2016). These profiles suggest that the small RNA profiles of viruses vary in different hosts, possibly due to variations in dicer isoforms processing viral dsRNAs among host species. Additionally, the vsiRNA ratio (the number of vsiRNA reads per virus transcript) varies among viruses in SBPH (Figure 6). Low-abundance viruses infecting insects can trigger robust host RNA interference responses and result in high-abundance vsiRNAs (Figure 6B). This effect may be attributed to the activity of viral suppressors of RNAi (VSRs), or differences in how viruses are targeted by insect immune mechanisms (Aguiar et al., 2015; Gammon & Mello, 2015; Webster et al., 2015). In addition to RNA interference, insects employ other mechanisms, such as Toll and IMD pathways, apoptosis, and autophagy, to control viral infections (Olmo et al., 2019). Therefore, studying the interactions and trade-offs between viruses and insects, as well as among different viruses, can enhance our understanding of the long-term adaptation of viruses to their hosts.

Virus transmission between host species is universal and underlies disease emergence (French et al., 2023). Host switching events occur readily between closely related host species, with the host genetic background considered a critical factor in shaping the insect virome (Huang et al., 2021; Longdon et al., 2014; Y. Xu et al., 2022). The three rice planthopper species used in this study share same habitat and many ecological characteristics, facilitating the possibility of cross-species transmission for these viruses. In this study, we demonstrated the ability of certain SBPH ISVs to infect other planthopper species through a microinjection approach (Figure 7), overcoming the pre-entry barrier. This suggests that similar virus-cell interactions occur during viral replication of certain viruses within two planthopper species. However, the presence of replication and assembly barriers limits the expansion of host range for most SBPH ISVs, indicating a relatively conservative phylosymbiosis between planthopper hosts and their viruses. Notably, for zoonotic viruses, host switching and spillover events have led to significant disease outbreaks in the past. Examples include the spillover of the Ebola virus from bats to humans and the recent COVID-19 pandemic, believed to have originated from a spillover event involving bats and an intermediate host (Ruiz-Aravena et al., 2022). Therefore, it is important to evaluate the transmission ability of viruses among different phytophagous insect hosts in future studies.

## Materials and Methods

### RNA sequencing (RNA-seq) Libraries construction

To investigate RNA viruses in SBPH, approximately 68 RNA-seq datasets were retrieved from the NCBI SRA repository. The dataset with the largest total number of bases was selected when there were several biological replicates available. As a result, a total of 28 typical high-quality SRA datasets were chosen representing at least 7 different lab populations from different universities and institutions in China. Meanwhile, lab-reared samples maintained in our phytotron in Ningbo University and field samples obtained from 6 rice-growing areas in China (Hangzhou, Jurong, Xinxiang, Xinzhou, Dalian and Shenyang), were collected to generate RNA-seq libraries. Total RNAs were extracted using 20-30 SBPHs comprised of different development stages from each sample. RNA samples were then used for Illumina high throughput sequencing (transcriptome). Paired-end (150 bp) sequencing of the RNA library was performed on the Illumina HiSeq 4000 platform (Illumina, USA) by Novogene (Tianjin, China). The transcriptome raw reads of these samples were deposited in Nature Microbiology Data Center (NMDC). A total of 39 transcriptome datasets were generated and the abbreviations and detail information are provided in Supplementary Table 1.

### Dataset reassembly and RNA virome discovery

Quality assessment were conducted for the sequencing reads of the 39 selected transcriptome datasets using FastQC and Trimmomatic. The filtered reads were reassembled/assembles de novo using the two assembler software packages Trinity and metaSPAdes with default parameters. The assembled contigs were compared with the NCBI viral RefSeq database using diamond Blastx. Since most of the datasets were retrieved from public databases, strict criteria were used for the identification of putative novel viruses in each dataset. Firstly, the diamond BlastX was set an E-value cutoff of 1×10^-20^. Secondly, the viral homology contigs had to meet a minimal coverage and length criteria of 20× and 500 bp, respectively, and they had to contain complete open reading frames (ORF) of predicted viral RNA-dependent RNA polymerase (RdRP). Thirdly, the viral homology contigs needed to be confirmed by both of the assemblers, Trinity and metaSPAdes. Fourthly, to eliminate false positives, the regions of the candidate viral-like contigs matched to the reference virus were further compared with the NCBI nucleotide and non-redundant protein databases. Sequences of all identified novel viruses from this study have been deposited in NMDC. Viral abundance was assessed using the number transcripts per million (TPM) within each library after the removal of rRNA reads.

### Virus genome annotation and phylogenetic analysis

The newly identified viral contigs were annotated with InterPro database (Mitchell et al., 2019). Conserved RdRP regions of the discovered viruses, together with RdRP protein sequences of reference viruses, were used for phylogenetic analysis. The RdRP sequences were aligned with MAFFT, and ambiguously aligned regions were then trimmed by Gblock. The best-fit model of amino acid substitution was evaluated by ModelTest-NG. Maximum likelihood (ML) trees were constructed using RAxML-NG with 1000 bootstrap replications.

### Virus names

Viruses identified in this work were named following these criteria: (i) The first part of the name is the latin name of host insect, *Laodelphax striatellus*; (ii) The second part of the name identifies is the virus taxonomic group. If the virus is assigned clearly to the level of “Genus”, the genus name is used. Additionally, if the virus can be only assigned to “Family” or “Order”, then the “prefix-like virus” is used. (iii) The third part of the name is a sequential number.

### Correlation for the RNA virome composition of SBPH with various development stages and tissues of the host insects

To investigate the relative temporal and spatial expression of ISVs in SBPH, samples from different development stages and tissues of adult insects were collected for RNA-seq. For viral temporal expression detection, SBPH eggs andnymphs of 1st-5th instar, as well as female and male adults at 0-24h, 2 day, 4 days, 6 days and 8 days post eclosion, were collected. Each sample comprised of 5-8 insects and two replicates were conducted. For viral spatial expression detection, tissue samples from intestines (Gut), salivary glands (SG), fat bodies (FB), male reproductive systems (MR), female reproductive systems (FR) and residues (Re) were dissected from the SBPH lab cultures. All the collected samples were conducted for RNA extraction and transcriptome RNA-seq. The viral abundance was subsequently compared in various tissues and development stages of SBPHs.

### Small RNA sequencing and analysis

The cDNA libraries were prepared using the Illumina TruSeq Small RNA Sample Preparation Kit (Illumina, CA, USA), and sRNA sequencing was performed on an Illumina HiSeq 2500 by Novogene (Tianjin, China). The raw sRNA reads were firstly treated to remove the adaptor, low quality, and junk sequences as described previously. The clean sRNA reads of the length 18- to 30-nt were extracted using the FASTX-Toolkit (http://hannonlab.cshl.edu/fastx_toolkit) and were mapped to the identified viral contigs using Bowtie software with perfect match (i.e. allowing zero mathch). Downstream analyses were performed using custom perl scripts and Linux shell bash scripts.

### Cross-species Ability Assessment of ISVs in the three rice planthoppers

To evaluate the infectivity of SBPH-ISVs to WBPH and BPH, the SBPH-ISVs inoculum were prepared from the SBPH populations as described previously (Huang et al., 2021). A total of 10 adult planthopper SBPHs were surface-sterilized with 70% ethanol and then rinsed with sterile distilled water. The sterilized insects were homogenized in 150 μl phosphate-buffered saline solutions and centrifuged. The supernatant was conducted for ISV detection using RT-PCR and then used as the inoculum microinjected into individual WBPH and BPH planthoppers. The injected insects were maintained on healthy rice seedlings and were collected at 0 and 6 days post-injection (DPI) for ISVs detection. Virus primers used for RT-qPCR were listed in Table S2, and actin gene of SBPH was served as the control. Relative transcript levels of viruses were calculated using the 2^-△△CT^ method. Additionally, the total RNA from microinjected insects at 6 DPI was sent to Novogene for small RNA sequencing.

## Data availability

The raw reads of RNA-seq generated in this study were deposited in NCBI SRA database and the accession numbers were listed in Supplementary Table S2.

## Acknowledgments

This work was supported by the National Natural Science Foundation of China (U20A2036, 32270146), the National Key Research and Development Plan in the 14^th^ five-year plan (2021YFD1401100: H.J.H. and C.X.Z.), and K.C. Wong Magna Fund in Ningbo University.

## Notes

### Competing Interest Statement

The authors have declared no competing interest.

## References

Aguiar, E. R. G. R., Olmo, R. P., & Marques, J. T. (2016). Virus-derived small RNAs: Molecular footprints of host–pathogen interactions. WIREs RNA, 7(6), 824–837. 10.1002/wrna.1361

Aguiar, E. R. G. R., Olmo, R. P., Paro, S., Ferreira, F. V., de Faria, I. J. da S., Todjro, Y. M. H., Lobo, F. P., Kroon, E. G., Meignin, C., Gatherer, D., Imler, J.-L., & Marques, J. T. (2015). Sequence-independent characterization of viruses based on the pattern of viral small RNAs produced by the host. Nucleic Acids Research, 43(13), 6191–6206. 10.1093/nar/gkv587

Ammar, E.-D., & Nault, L. R. (2002). Virus transmission by leafhoppers, planthoppers and treehoppers (auchenorrhyncha, homoptera). In Advances in Botanical Research (Vol. 36, pp. 141–167). Elsevier. 10.1016/S0065-2296(02)36062-2

Baidaliuk, A., Miot, E. F., Lequime, S., Moltini-Conclois, I., Delaigue, F., Dabo, S., Dickson, L. B., Aubry, F., Merkling, S. H., Cao-Lormeau, V.-M., & Lambrechts, L. (2019). Cell-Fusing Agent Virus Reduces Arbovirus Dissemination in Aedes aegypti Mosquitoes *In Vivo*. Journal of Virology, 93(18), e00705–19. 10.1128/JVI.00705-19

Bonning, B. C. (2019). The Insect Virome: Opportunities and Challenges. In Insect Molecular Virology: Advances and Emerging Trends. Caister Academic Press. 10.21775/9781912530083.01

Bonning, B. C., & Saleh, M.-C. (2021). The Interplay Between Viruses and RNAi Pathways in Insects. Annual Review of Entomology, 66(1), 61–79. 10.1146/annurev-ento-033020-090410

Cao, Q., Xu, W.-Y., Gao, Q., Jiang, Z.-H., Liu, S.-Y., Fang, X.-D., Gao, D.-M., Wang, Y., & Wang, X.-B. (2018). Transmission Characteristics of Barley Yellow Striate Mosaic Virus in Its Planthopper Vector Laodelphax striatellus. Frontiers in Microbiology, 9, 1419. 10.3389/fmicb.2018.01419

Chandler, J. A., Liu, R. M., & Bennett, S. N. (2015). RNA shotgun metagenomic sequencing of northern California (USA) mosquitoes uncovers viruses, bacteria, and fungi. Frontiers in Microbiology, 6. https://www.frontiersin.org/articles/10.3389/fmicb.2015.00185

Chen, D.-S., Yang, S.-X., Ding, X.-L., Zhang, Y.-K., & Hong, X.-Y. (2015). Infection Rate Assay by Nested PCR and the Phylogenetic Analysis of Himetobi P Virus in the Main Pests of Rice-Wheat Cropping Systems. Journal of Economic Entomology, 108(3), 1304–1312. 10.1093/jee/tov001

Chiapello, M., Bosco, L., Ciuffo, M., Ottati, S., Salem, N., Rosa, C., Tavella, L., & Turina, M. (2021). Complexity and Local Specificity of the Virome Associated with Tospovirus-Transmitting Thrips Species. Journal of Virology, 95(21), e00597–21. 10.1128/JVI.00597-21

Dance, A. (2021). The incredible diversity of viruses. Nature, 4.

de Almeida, J. P., Aguiar, E. R., Armache, J. N., Olmo, R. P., & Marques, J. T. (2021). The virome of vector mosquitoes. Current Opinion in Virology, 49, 7–12. 10.1016/j.coviro.2021.04.002

Di Paola, N., Dheilly, N. M., Junglen, S., Paraskevopoulou, S., Postler, T. S., Shi, M., & Kuhn, J. H. (2022). *Jingchuvirales*: A New Taxonomical Framework for a Rapidly Expanding Order of Unusual Monjiviricete Viruses Broadly Distributed among Arthropod Subphyla. Applied and Environmental Microbiology, 88(6), e01954–21. 10.1128/aem.01954-21

Dolja, V. V., Krupovic, M., & Koonin, E. V. (2020). Deep Roots and Splendid Boughs of the Global Plant Virome. Annual Review of Phytopathology, 58(1), 23–53. 10.1146/annurev-phyto-030320-041346

Feng, Y., Gou, Q., Yang, W., Wu, W., Wang, J., Holmes, E. C., Liang, G., & Shi, M. (2022). A time-series meta-transcriptomic analysis reveals the seasonal, host, and gender structure of mosquito viromes. Virus Evolution, 8(1), veac006. 10.1093/ve/veac006

French, R. K., Anderson, S. H., Cain, K. E., Greene, T. C., Minor, M., Miskelly, C. M., Montoya, J. M., Wille, M., Muller, C. G., Taylor, M. W., Digby, A., Kākāpō Recovery Team, Crane, J., Davitt, G., Eason, D., Hedman, P., Jeynes, B., Latimer, S., Little, S., Holmes, E. C. (2023). Host phylogeny shapes viral transmission networks in an island ecosystem. Nature Ecology & Evolution. 10.1038/s41559-023-02192-9

Gammon, D. B., & Mello, C. C. (2015). RNA interference-mediated antiviral defense in insects. Current Opinion in Insect Science, 8, 111–120. 10.1016/j.cois.2015.01.006

Goenaga, S., Kenney, J., Duggal, N., Delorey, M., Ebel, G., Zhang, B., Levis, S., Enria, D., & Brault, A. (2015). Potential for Co-Infection of a Mosquito-Specific Flavivirus, Nhumirim Virus, to Block West Nile Virus Transmission in Mosquitoes. Viruses, 7(11), 5801–5812. 10.3390/v7112911

Gong, W., Liu, H., Zhu, X., Zhao, S., Cheng, J., Zhu, H., Zhong, J., & Zhou, Q. (2021). Molecular characterization of a novel fusarivirus infecting the plant-pathogenic fungus Alternaria solani. Archives of Virology, 166(7), 2063–2067. 10.1007/s00705-021-05105-y

GuY, P. L., Tofiyama, S., & Fuji, S. (1988). Occurrence of a Picorna-like Virus in Planthopper Species and Its Transmission in Laodelphax striatellus. Journal of Invertebrate Pathology, 4.

Hillman, B. I., & Cai, G. (2013). Chapter Six - The Family Narnaviridae: Simplest of RNA Viruses. In S. A. Ghabrial (Ed.), Advances in Virus Research (Vol. 86, pp. 149–176). Academic Press. 10.1016/B978-0-12-394315-6.00006-4

Huang, H.-J., Ye, Z.-X., Wang, X., Yan, X.-T., Zhang, Y., He, Y.-J., Qi, Y.-H., Zhang, X.-D., Zhuo, J.-C., Lu, G., Lu, J.-B., Mao, Q.-Z., Sun, Z.-T., Yan, F., Chen, J.-P., Zhang, C.-X., & Li, J.-M. (2021). Diversity and infectivity of the RNA virome among different cryptic species of an agriculturally important insect vector: Whitefly Bemisia tabaci. Npj Biofilms and Microbiomes, 7(1), 43. 10.1038/s41522-021-00216-5

Jia, W., Wang, F., Li, J., Chang, X., Yang, Y., Yao, H., Bao, Y., Song, Q., & Ye, G. (2021). A Novel Iflavirus Was Discovered in Green Rice Leafhopper Nephotettix cincticeps and Its Proliferation Was Inhibited by Infection of Rice Dwarf Virus. Frontiers in Microbiology, 11, 621141. 10.3389/fmicb.2020.621141

Koonin, E. V., Krupovic, M., & Dolja, V. V. (2022). The global virome: How much diversity and how many independent origins? Environmental Microbiology, 1462–2920.16207. 10.1111/1462-2920.16207

Lacey, L. A., Grzywacz, D., Shapiro-Ilan, D. I., Frutos, R., Brownbridge, M., & Goettel, M. S. (2015). Insect pathogens as biological control agents: Back to the future. Journal of Invertebrate Pathology, 132, 1–41. 10.1016/j.jip.2015.07.009

Lan, H., Wang, H., Chen, Q., Chen, H., Jia, D., Mao, Q., & Wei, T. (2016). Small interfering RNA pathway modulates persistent infection of a plant virus in its insect vector. Scientific Reports, 6(1), 20699. 10.1038/srep20699

Li, J., Andika, I. B., Shen, J., Lv, Y., Ji, Y., Sun, L., & Chen, J. (2013). Characterization of Rice Black-Streaked Dwarf Virus- and Rice Stripe Virus-Derived siRNAs in Singly and Doubly Infected Insect Vector Laodelphax striatellus. PLoS ONE, 8(6), e66007. 10.1371/journal.pone.0066007

Li, J., Andika, I. B., Zhou, Y., Shen, J., Sun, Z., Wang, X., Sun, L., & Chen, J. (2014). Unusual characteristics of dicistrovirus-derived small RNAs in the small brown planthopper, Laodelphax striatellus. Journal of General Virology, 95(3), 712–718. 10.1099/vir.0.059626-0

Li, N., Li, C., Hu, T., Li, J., Zhou, H., Ji, J., Wu, J., Kang, W., Holmes, E. C., Shi, W., & Xu, S. (2023). Nationwide genomic surveillance reveals the prevalence and evolution of honeybee viruses in China. Microbiome, 11(1), 6. 10.1186/s40168-022-01446-1

Longdon, B., Brockhurst, M. A., Russell, C. A., Welch, J. J., & Jiggins, F. M. (2014). The Evolution and Genetics of Virus Host Shifts. PLoS Pathogens, 10(11), e1004395. 10.1371/journal.ppat.1004395

Lu, G., Zhang, X.-D., Xu, Z.-T., Ye, Z.-X., Zhang, Y., Chen, J.-P., Zhang, C.-X., & Li, J.-M. (2022a). Complete sequence and genetic characterization of a novel insect-specific reovirus discovered from Laodelphax striatellus. Virology, 570, 117–122. 10.1016/j.virol.2022.03.011

Lu, G., Zhang, X.-D., Xu, Z.-T., Ye, Z.-X., Zhang, Y., Chen, J.-P., Zhang, C.-X., & Li, J.-M. (2022b). Complete sequence and genetic characterization of a novel insect-specific reovirus discovered from Laodelphax striatellus. Virology, 570, 117–122. 10.1016/j.virol.2022.03.011

Mitchell, A. L., Attwood, T. K., Babbitt, P. C., Blum, M., Bork, P., Bridge, A., Brown, S. D., Chang, H.-Y., El-Gebali, S., Fraser, M. I., Gough, J., Haft, D. R., Huang, H., Letunic, I., Lopez, R., Luciani, A., Madeira, F., Marchler-Bauer, A., Mi, H., Finn, R. D. (2019). InterPro in 2019: Improving coverage, classification and access to protein sequence annotations. Nucleic Acids Research, 47(D1), D351–D360. 10.1093/nar/gky1100

Nault, L. R., & Ammar, E. D. (1989). Leafhopper and Planthopper Transmission of Plant Viruses. Annual Review of Entomology, 34:503–29.

Nouri, S., Matsumura, E. E., Kuo, Y.-W., & Falk, B. W. (2018). Insect-specific viruses: From discovery to potential translational applications. Current Opinion in Virology, 33, 33–41. 10.1016/j.coviro.2018.07.006

Öhlund, P., Lundén, H., & Blomström, A.-L. (2019). Insect-specific virus evolution and potential effects on vector competence. Virus Genes, 55(2), 127–137. 10.1007/s11262-018-01629-9

Olmo, R. P., Martins, N. E., Aguiar, E. R. G. R., Marques, J. T., & Imler, J.-L. (2019). The insect reservoir of biodiversity for viruses and for antiviral mechanisms. Anais Da Academia Brasileira de Ciências, 91. 10.1590/0001-3765201920190122

Olmo, R. P., Todjro, Y. M. H., Aguiar, E. R. G. R., de Almeida, J. P. P., Ferreira, F. V., Armache, J. N., de Faria, I. J. S., Ferreira, A. G. A., Amadou, S. C. G., Silva, A. T. S., de Souza, K. P. R., Vilela, A. P. P., Babarit, A., Tan, C. H., Diallo, M., Gaye, A., Paupy, C., Obame-Nkoghe, J., Visser, T. M., Marques, J. T. (2023). Mosquito vector competence for dengue is modulated by insect-specific viruses. Nature Microbiology, 8(1), 135–149. 10.1038/s41564-022-01289-4

Patterson, E. I., Villinger, J., Muthoni, J. N., Dobel-Ober, L., & Hughes, G. L. (2020). Exploiting insect-specific viruses as a novel strategy to control vector-borne disease. Current Opinion in Insect Science, 39, 50–56. 10.1016/j.cois.2020.02.005

Roossinck, M. J. (2019). Evolutionary and ecological links between plant and fungal viruses. New Phytologist, 221(1), 86–92. 10.1111/nph.15364

Ruiz-Aravena, M., McKee, C., Gamble, A., Lunn, T., Morris, A., Snedden, C. E., Yinda, C. K., Port, J. R., Buchholz, D. W., Yeo, Y. Y., Faust, C., Jax, E., Dee, L., Jones, D. N., Kessler, M. K., Falvo, C., Crowley, D., Bharti, N., Brook, C. E., Plowright, R. K. (2022). Ecology, evolution and spillover of coronaviruses from bats. Nature Reviews Microbiology, 20(5), 299–314. 10.1038/s41579-021-00652-2

Shi, M., Lin, X.-D., Tian, J.-H., Chen, L.-J., Chen, X., Li, C.-X., Qin, X.-C., Li, J., Cao, J.-P., Eden, J.-S., Buchmann, J., Wang, W., Xu, J., Holmes, E. C., & Zhang, Y.-Z. (2016). Redefining the invertebrate RNA virosphere. Nature, 540(7634), 539–543. 10.1038/nature20167

Sõmera, M., Fargette, D., Hébrard, E., Sarmiento, C., & Ictv Report Consortium, null. (2021). ICTV Virus Taxonomy Profile: Solemoviridae 2021. The Journal of General Virology, 102(12), 001707. 10.1099/jgv.0.001707

Wang, F., Fang, Q., Wang, B., Yan, Z., Hong, J., Bao, Y., Kuhn, J. H., Werren, J. H., Song, Q., & Ye, G. (2017). A novel negative-stranded RNA virus mediates sex ratio in its parasitoid host. PLOS Pathogens, 13(3), e1006201. 10.1371/journal.ppat.1006201

Webster, C. L., Waldron, F. M., Robertson, S., Crowson, D., Ferrari, G., Quintana, J. F., Brouqui, J.-M., Bayne, E. H., Longdon, B., Buck, A. H., Lazzaro, B. P., Akorli, J., Haddrill, P. R., & Obbard, D. J. (2015). The Discovery, Distribution, and Evolution of Viruses Associated with Drosophila melanogaster. PLOS Biology, 13(7), e1002210. 10.1371/journal.pbio.1002210

Wu, N., Zhang, P., Liu, W., Cao, M., Massart, S., & Wang, X. (2019). Complete genome sequence and characterization of a new iflavirus from the small brown planthopper (Laodelphax striatellus). Virus Research, 272, 197651. 10.1016/j.virusres.2019.197651

Xu, P., Yang, L., Yang, X., Li, T., Graham, R. I., Wu, K., & Wilson, K. (2020). Novel partiti-like viruses are conditional mutualistic symbionts in their normal lepidopteran host, African armyworm, but parasitic in a novel host, Fall armyworm. PLOS Pathogens, 16(6), e1008467. 10.1371/journal.ppat.1008467

Xu, Y., Jiang, J., Lin, X., Shi, W., & Cao, C. (2022). Identification of diverse viruses associated with grasshoppers unveils the parallel relationship between host phylogeny and virome composition. Virus Evolution, 8(2), veac057. 10.1093/ve/veac057

