## Supplemental Figure 1, Supplementary Table 1, Supplementary Table 2 for "Diversity and adaptability of RNA viruses in rice planthopper, *Laodelphax striatellus*"

Sampling locations of small brown planthoppers (SBPH).

**Supplementary Table 1. SBPH datasets used in this study.**

**Supplementary Table 2. RNA virome identified in SBPH in this study.**

**Supplemental Figure 1**

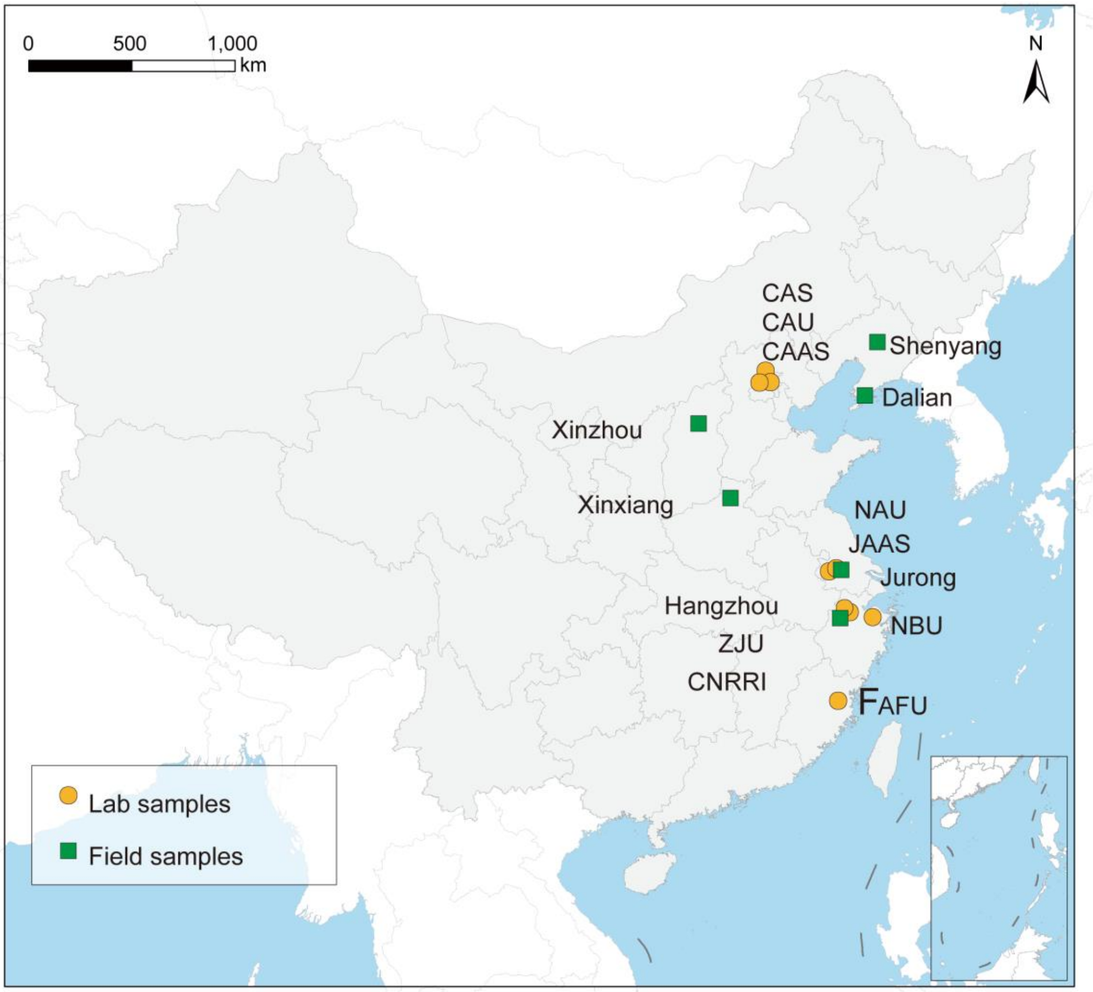

#### Supplementary Table 1. SBPH datasets used in this study

| Library | Run Accession Number | Brief Description | Total Base (Gb) | N50 (nt) | BioProject Accession | Submitter | Location |
|---|---|---|---|---|---|---|---|
| FAFU-lab1 | SRR7058093 | RSV infected SBPH | 6.1 | | PRJNA451497 | Fujian Agriculture and Forestry University | Fuzhou, China |
| ZJU-lab1 | SRR10090680 | SBPH(gut) feed on wheat | 7.9 | 2297 | PRJNA564687 | Zhejiang University | Hangzhou, China |
| ZJU-lab2 | SRR10090682 | SBPH(gut) feed on rice | 8.4 | 1918 | | | |
| ZJU-lab3 | SRR10090691 | SBPH(gut) transferred from rice to wheat | 8.1 | 2324 | | | |
| ZJU- lab4 | SRR4020768 | SBPH adult | 4.7 | 886 | PRJNA338373 | | |
| CNRRI-lab1 | SRR11729951 | SBPH adult | 10.4 | 1704 | PRJNA629998 | China National Rice Research Institute | |
| NAU-lab1 | SRR4088002 | Wolbachia-infected SBPH | 27.4 | 3032 | PRJNA340357 | Nanjing Agricultural University | Nanjing, China |
| NAU-lab2 | SRR4088021 | Wolbachia-uninfected SBPH | 33.6 | 2633 | | | |
| NAU-lab3 | SRR4075582 | Wolbachia-infected L. striatellus1-1 | 14.8 | 3012 | PRJNA340307 | | |
| NAU-lab4 | SRR4075604 | Wolbachia-uninfected SBPH | 18.1 | | | | |
| NAU-lab5 | SRR4087171 | Wolbachia-infected SBPH | 14.8 | 3071 | | | |
| NAU-lab6 | SRR941775 | Mix different development stages for SBPH | 6.0 | 782 | PRJNA212544 | | |
| JAAS-lab1 | SRR12076774 | RBSDV infection in SBPH midgut | 15.0 | 1375 | | | |
| JAAS-lab2 | SRR12076777 | RBSDV free in SBPH midgut | 15.8 | 1235 | | | |
| CAS-lab1 | SRR5816374 | SBPH gonad | 9.6 | 2321 | PRJNA393384 | Institute of Zoology, Chinese Academy of Sciences | Beijing,China |
| CAS-lab2 | SRR5816375 | SBPH brain | 11.5 | 2340 | | | |
| CAS-lab3 | SRR5816380 | SBPH adult | 7.3 | 2603 | | | |
| CAS-lab4 | SRR5816381 | SBPH larva | 7.0 | 2615 | | | |

| Library | Run Accession Number | Brief Description | Total Base (Gb) | N50 (nt) | BioProject Accession | Submitter | Location |
|---|---|---|---|---|---|---|---|
| CAS-lab5 | SRR5816382 | SBPH fat body | 9.0 | 2479 | | | |
| CAS-lab6 | SRR5816383 | SBPH egg | 7.2 | 2681 | | | |
| CAS-lab7 | SRR1614218 | SBPH naive Salivary gland | 4.6 | 1426 | PRJNA263985 | | |
| CAS-lab8 | SRR1617617 | SBPH RSV virulent gut | 5.5 | 2244 | | | |
| CAS-lab9 | SRR1617623 | SBPH naive gut | 4.6 | 2091 | | | |
| CAS-lab10 | SRR1619428 | SBPH RSV virulent Salivary gland | 4.7 | 926 | | | |
| CAU-lab1 | SRR5814450 | SBPH BYSMV virulent hindgut | 15.1 | 1461 | PRJNA393404 | China Agricultural University | |
| CAU-lab2 | SRR5814453 | SBPH naive hindgut | 12.7 | 2002 | | | |
| CAAS-lab1 | SRR7091178 | 48 hours SBPH RSV virulent ovary | 11.2 | 2292 | PRJNA450301 | Chinese Academy of Agricultural Sciences | |
| CAAS-lab2 | SRR7091179 | 24 hours SBPH non-virulent ovary | 9.9 | 2194 | | | |
| NBU-lab1 | | SBPH adults of lab population | | | | Ningbo University | Ningbo, China |
| NBU-lab2 | | SBPH adults of lab population Ribosomal RNA-depleted libraries | | | | | |
| NBU-HZ1 | | Offspring of field population | | | | | Hangzhou, China |
| NBU-HZ2 | | Offspring of field population | | | | | |
| NBU-Jurong | | SBPH adults collected from field | | | | | Jurong, China |
| NBU-Xinxiang | | SBPH adults collected from field | | | | | Xinxiang, China |
| NBU-Xinzhou | | SBPH adults collected from field | | | | | Xinzhou, China |
| NUB-Dalian | | SBPH adults collected from field | | | | | Daliang, China |
| NBU-Shenyang1 | | SBPH adults collected from field | | | | | Shenyang, China |
| NBU-Shenyang2 | | SBPH adults collected from field | | | | | |

### Supplementary Table 2. RNA virome identified in SBPH in this study

| | Virus names | | NCBI Accession | Length (nt) | Coverage | E-value | Homology virus (genome size, accession) | Protein Identities | Order | Family | Genus |
|---|---|---|---|---|---|---|---|---|---|---|---|
| dsRNA viruses | Laodelphax striatellus alphafusarivirus 1 (LSAfV1) | | NMDCN0001DK1 | 5601 | | | Penicillium roqueforti ssRNA mycovirus 1 （6002 nt, NC_024699.1） | | *Durnavirales* | *Fusariviridae* | *Alphafusarivirus* |
| | Laodelphax striatellus partiti-like virus 1 (LSPalV1) | | NMDCN0001DJI | 1480 | | 7e-159 | Hubei partiti-like virus 46 (1444 nt, KX884130.1) [1] | 51.66% | | *Partitiviridae* | Unassigned |
| | Laodelphax striatellus reovirus (LSRV) | S1 | OM249649 | 4436 | / | / | / | / | *Reovirales* | *Spinareoviridae* | *Fijivirus* |
| | | S2 | OM249650 | 3643 | / | / | | / | | | |
| | | S3 | OM249651 | 3804 | / | / | | / | | | |
| | | S4 | OM249652 | 3628 | / | / | | / | | | |
| | | S5 | OM249653 | 2872 | / | / | | / | | | |
| | | S6 | OM249654 | 2825 | / | / | | / | | | |
| | | S7 | OM249655 | 1931 | / | / | | / | | | |
| | | S8 | OM249656 | 1837 | / | / | | / | | | |
| | | S9 | OM249657 | 1656 | / | / | | / | | | |
| | | S10 | OM249658 | 1575 | / | / | | / | | | |
| +ssRNA virus | Laodelphax striatellus permutotetra-like virus 1 (LSPelV1) | | NMDCN0001DJJ | 4641 | | 0.0 | Daeseongdong virus 2 (4742 nt, NC_028489.1) [2] | 46.94% | / | *Permutotetraviridae* | Unassigned |
| | Laodelphax striatellus flavi-like virus 1 (LSFlV1) | | NMDCN0001DJN | 22292 | | | Fushun laodelphax striatellus flavivirus 1 (22293 nt, MZ210009.1) | 99.99% | *Amarillovirales* | *Flaviviridae* | Unassigned |
| | Laodelphax striatellus Botourmia-like virus 1 (LSBolV1) | | NMDCNO001DJV | 2478 | | 0.0 | Erysiphe necator associated ourmia-like virus 14 ( 2441 nt, MN611542.1) | 72.43% | *Ourlivirales* | *Botourmiaviridae* | Unassigned |
| | Laodelphax striatellus Botourmia-like virus 2 (LSBolV2) | | NMDCNO001DKO | 1921 | | 0.0 | Plasmopara viticola lesion associated ourmia-like virus 79 (2101 nt, MN532666.1) [8] | 71.23% | | | Unassigned |
| | Laodelphax striatellus iflavirus 2 (LSIfV2) | | NMDCN0001DJM | 6041 | | 0.0 | Bemisia tabaci iflavirus 1 (7812 nt, MW256671.1) [4] | 35.71% | *Picornavirales* | *Iflaviridae* | *Iflavirus* |

| | | | | | | | | | | |
|---|---|---|---|---|---|---|---|---|---|---|
| | Laodelphax striatellus iflavirus 1 (LSIV1) | MG815140.1[11] | 10831 | / | / | / | / | | | |
| | Laodelphax striatellus picorna-like virus 2 (LSPV2) | NC_025788.1 | 10889 | / | / | / | / | | | |
| | Laodelphax striatella honeydew virus 1 (LSHV1) | NC_023627.1 | 10929 | / | / | / | / | | | |
| | Himetobi P virus (HiPV) | KF280585.1[10] | 9275 | / | / | / | / | | *Dicistroviridae* | *Triatovirus* |
| | Laodelphax striatellus sobemovirus 1 (LSSoV1) | NMDCN0001DJQ | 3458 | | 4e-80 | Jilin luteo/like virus 2 (2096 nt, MW239178.1) | 33.27% | *Sobelivirales* | *Solemoviridae* | *Sobemovirus* |
| | Laodelphax striatellus sobemovirus 2 (LSSoV2) | NMDCN0001DJR | 3267 | | 4e-119 | Norway luteo/like virus 1 (3250 nt, MF141066.1[6]) | 49.52% | | | |
| | Laodelphax striatellus sobemovirus 3 (LSSoV3) | NMDCN0001DJO | 3152 | | 2e/86 | Amygdalus persica sobemo/like virus (3255 nt, MN831439.1[5]) | 42.15% | | | |
| | Laodelphax striatellus sobemovirus 4 (LSSoV4) | NMDCNO001DJP | 3403 | | 8e-71 | Atrato Sobemo/like virus 5 (2733nt, MN661099.1) | 34.72% | | | |
| | Laodelphax striatellus narna-like virus 1 (LSNaV1) | NMDCN0001DJK | 2844 | | 0.0 | Serbia narna/like virus 3 (2685 nt, MT822185.1) [3] | 58.04% | *Wolframvirales* | *Narnaviridae* | Unassigned |
| | Laodelphax striatellus narna-like virus 2 (LSNaV2) | NMDCN0001DJL | 3447 | | 0.0 | Wenling narna/like virus 6 (3331 nt, KX883609.1) [1] | 35.88% | | | |
| | Laodelphax striatellus bunya-like virus 1 (LSBulV1) | NMDCN0001DJS | 7003 | | 0.0 | Fushun phasmavirus 2 (2888 nt, MZ210012.1) | 99.72% | *Bunyavirales* | Unassigned | Unassigned |
| | Laodelphax striatellus Mobuvirus 1 (LSMoV1) | NMDCN0001DJU | 6594 | | 0.0 | Sanya nilaparvata lugens phenuivirus 1 (6957 nt, MZ209823.1) | 44.51% | | *Phenuiviridae* | *Mobuvirus* |
| | Laodelphax striatellus phenui-like virus 1 (LSPhlV1) | NMDCN0001DJT | 7454 | | 0.0 | Aphis citricidus bunyavirus (7037 nt, MN163034.1) [7] | 31.58% | | | Unassigned |
| -ssRNA virus | Laodelphax striatellus ollusvirus 1 (LSOlV1) | NMDCN0001DK2 | 15373 | | 0.0 | Hymenopteran chu/related virus OKIAV125 (7530 nt, MW039255.1[9]) | 39.22% | *Jingchuvirales* | *Aliusviridae* | *Ollusvirus* |

[1] SHI M, LIN X/D, TIAN J/H, et al. Redefining the invertebrate RNA virosphere.[J]. Nature, England: 2016, 540(7634): 539–543. DOI:10.1038/nature20167.
[2] HANG J, KLEIN T A, KIM H/C, et al. Genome Sequences of Five Arboviruses in Field/Captured Mosquitoes in a Unique Rural Environment of South Korea[J]. Genome Announcements, 2016, 4(1): e01644/15. DOI:10.1128/genomeA.01644/15.
[3] STANOJEVIĆ M, LI K, STAMENKOVIĆ G, et al. Depicting the RNA Virome of Hematophagous Arthropods from Belgrade, Serbia[J]. Viruses, 2020, 12(9): E975. DOI:10.3390/v12090975.
[4] HUANG H/J, YE Z/X, WANG X, et al. Diversity and infectivity of the RNA virome among different cryptic species of an agriculturally important insect vector: whitefly Bemisia tabaci[J]. npj Biofilms and Microbiomes, Nature Publishing Group, 2021, 7(1): 1–15. DOI:10.1038/s41522/021/00216/5.
[5] YANG S, SHAN T, WANG Y, et al. Virome of riverside phytocommunity ecosystem of an ancient canal[R]. In Review, 2020. DOI:10.21203/rs.3.rs/25620/v1.
[6] PETTERSSON J H/O, SHI M, BOHLIN J, et al. Characterizing the virome of Ixodes ricinus ticks from northern Europe.[J]. Scientific reports, Nature Publishing Group, 2017, 7(1): 10870. DOI:10.1038/s41598/017/11439/y.
[7] ZHANG W, WU T, GUO M, et al. Characterization of a new bunyavirus and its derived small RNAs in the brown citrus aphid, Aphis citricidus[J]. Virus Genes, 2019, 55(4): 557–561. DOI:10.1007/s11262/019/01667/x.
[8] CHIAPELLO M, RODRÍGUEZ/ROMERO J, AYLLÓN M A, et al. Analysis of the virome associated to grapevine downy mildew lesions reveals new mycovirus lineages[J]. Virus Evolution, 2020, 6(2): veaa058. DOI:10.1093/ve/veaa058.
[9] KÄFER S, PARASKEVOPOULOU S, ZIRKEL F, et al. Re/assessing the diversity of negative strand RNA viruses in insects[J]. PLoS pathogens, 2019, 15(12): e1008224. DOI:10.1371/journal.ppat.1008224.
[10] XU Y, HUANG L, WANG Z, et al. Identification of Himetobi P virus in the small brown planthopper by deep sequencing and assembly of virus/derived small interfering RNAs[J]. Virus Research, 2014, 179: 235–240. DOI:10.1016/j.virusres.2013.11.004.
[11] WU N, ZHANG P, LIU W, et al. Complete genome sequence and characterization of a new iflavirus from the small brown planthopper (Laodelphax striatellus)[J]. Virus Research, 2019, 272: 197651. DOI:10.1016/j.virusres.2019.197651.
